# Methods for High-Throughput Drug Combination Screening and Synergy Scoring

**DOI:** 10.1101/051698

**Authors:** Liye He, Evgeny Kulesskiy, Jani Saarela, Laura Turunen, Krister Wennerberg, Tero Aittokallio, Jing Tang

## Abstract

Gene products or pathways that are aberrantly activated in cancer but not in normal tissue hold great promises for being effective and safe anticancer therapeutic targets. Many targeted drugs have entered clinical trials but so far showed limited efficacy mostly due to variability in treatment responses and often rapidly emerging resistance. Towards more effective treatment options, we will critically need multi-targeted drugs or drug combinations, which selectively inhibit the cancer cells and block distinct escape mechanisms for the cells to become resistant. Functional profiling of drug combinations requires careful experimental design and robust data analysis approaches. At the Institute for Molecular Medicine Finland (FIMM), we have developed an experimental-computational pipeline for high-throughput screening of drug combination effects in cancer cells. The integration of automated screening techniques with advanced synergy scoring tools allows for efficient and reliable detection of synergistic drug interactions within a specific window of concentrations, hence accelerating the identification of potential drug combinations for further confirmatory studies.

## 1. Introduction

A pressing challenge in the development of personalized cancer medicine is to understand how to make the most out of genomic information from a patient when evaluating treatment options. Over the past decade, there has been an extensive effort to sequence cancer genomes in large patient cohorts, sparking expectations to identify novel targets for more effective and selective treatment opportunities. These sequencing efforts have revealed a remarkable degree of genetic heterogeneity between and within tumors, which partly explains why the traditional ‘one-size-fits-all’ anticancer treatment strategies have produced many disappointing outcomes in clinical trials ***(1)***. On the other hand, functional studies using high-throughput drug screening have enabled linking cancer genomic vulnerabilities to targeted drug responses ***(2–4)***. However, complex genetic and epigenetic changes may lead to re-activation of multiple compensatory pathways and to emergence of treatment-resistant sub-populations (so-called cancer clonal evolution). Therefore, to reach effective and sustained clinical responses, one often needs multi-targeted drugs or drug combinations, which selectively inhibit multiple cancer driving sub-clones and other escape pathways of the cancer cells ***(5, 6)***. To facilitate the drug combination discovery, preclinical studies often rely on drug combination screening in cancer cell models as a starting point to prioritize the most potential hits for further experimental investigation and therapy optimization. Many of the existing drug combination studies, however, focus on conventional chemotherapeutic drugs tested on a panel of cell lines, for which the drug combination effects might not easily translate into treatment options in the clinic (see e.g. ***(7)***). Rather, cancer cells that are derived from patients have shown tremendous potential that could enable the rapid assessment of novel drugs or drug combinations at the individual level ***(8)***. To facilitate clinical translation, an Individualized Systems Medicine (ISM) drug combination platform has been established at FIMM that combines genomics, drug testing and computational tools to predict drug responses for individual cancer patients. The ISM platform has successfully been used to functionally profile primary leukemia, ovarian cancer and prostate cancer samples *ex vivo* so that the drug responses can be translated to the *in vivo* setting ***(9–12)***.

The advances in high-throughput drug combination screening has enabled the assaying of a large collection of chemical compounds, generating dynamic dose-response profiles that allow us to quantify the degree of drug-drug interactions at an unprecedented level. A drug interaction is usually classified as synergistic, antagonistic or non-interactive, based on the deviation of the observed drug combination response from the expected effect of non-interaction (the null hypothesis). To quantify the degree of drug synergy, several models have been proposed, such as those based on the Highest single agent model (HSA) ***(13)***, the Loewe additivity model (Loewe) ***(14)*** and the Bliss independence model (Bliss) ***(15)***. However, these existing drug synergy scoring models, together with their software implementations, were proposed initially for low-throughput experiments, with a limited number of drugs being combined with a fixed level of response, e.g. at their IC50 concentrations. For example, CompuSyn has become a popular tool to calculate a combination index (CI) using the Loewe additivity model ***(16)***. However, CompuSyn allows only for manual input of one drug combination at a time, which makes it less efficient for analyzing multiple drug combinations, particularly when the drug combinations are tested under various concentrations, in a so-called dose-response matrix design.

To facilitate the data analysis of high-throughput drug combination screens, more recent tools have been made available as R implementations (https://www.R-project.org). For example, mixlow is an R package, which utilizes a nonlinear mixed-effects model to calculate the CI ***(17)***. However, mixlow works only for an experimental design where the ratio of two drugs in a combination is fixed over all the tested concentrations. Therefore, it may not be directly applicable for a dose-response matrix design, where the ratios of two drugs vary. Another R package, called drc, provides an URSA (universal response surface approach) model, which is more suitable for dose-response matrix data ***(18)***. URSA extends the Loewe model by considering the response surfaces over all the tested concentrations. In contrast to the CI, which is defined at a fixed response level, the URSA model provides a summarized drug interaction score from the whole dose-response matrix. However, the URSA implementation in the drc package often leads to fitting errors when the dose responses fail to comply with the model assumptions. To evaluate the appropriateness of URSA, one needs to trace back to its underlying theoretical paper ***(19)***, which becomes rather technical for end users. The Bliss model has also been extended recently by incorporating the response surface concept, similar as in the URSA model, based on which a contour plot of a Bliss interaction index can be constructed ***(20)***. We have recently developed a response surface model, called Zero Interaction Potency (ZIP), which combines the Loewe and the Bliss models, and proposed a delta score to characterize the synergy landscape over the full dose-response matrix ***(21)***.

Here, we describe a seamless experimental-computational drug combination analysis pipeline that has been widely used in Finland and elsewhere to test and score effects of drug combinations in cancer cells ***(22–25)***. The pipeline includes both an experimental protocol for dose-response matrix drug combination assays, as well as computational tools to facilitate the plate design and synergy modeling. The pipeline is applicable not only to cancer cell lines but also to patient-derived cancer samples for individualized drug combination optimization. With the increasing size of our compound library, targeting all the known cancer survival pathways, the drug combination discovery is now possible towards more personalized anticancer treatment. We first describe the experimental protocol including a computer program, called FIMMcherry, which enables efficient production and visualization of combination assay plates, the output of which can be directly exported to the robotic system for automated dispensing. To address the lack of tailored software tools for high throughput drug combination scoring, we report here a new R package, synergyfinder, which provides efficient implementations for all the popular synergy scoring models, including HSA, Loewe, Bliss and ZIP. This implementation provides the lab users with more flexibility to explore their drug combination data. We expect that the use of synergyfinder will greatly improve the interpretation of the drug combination results and may eventually lead to the standardization of preclinical drug combination studies.

## 2. Materials

### 2.1. Cancer cells or patient-derived samples

1. Established cancer cell lines are purchased from multiple vendors.
2. Patient-derived samples are obtained with permission from Finnish biobanks, hospitals and clinical collaborators ***(2)***.

### 2.2 Compounds

The FIMM oncology collection contains both FDA/EMA-approved drugs as well as investigational compounds (Fig. 1). The collection is constantly evolving and the current FO4B version contains 525 compounds with concentrations ranging typically between 1-10,000 nM. For some compounds, the concentration range is adjusted upwards (e.g. platinum drugs, 100,000 nM) or downwards (e.g. rapalogs, 100 nM) to better match their relevant concentrations of bioactivity.

**Fig. 1.**
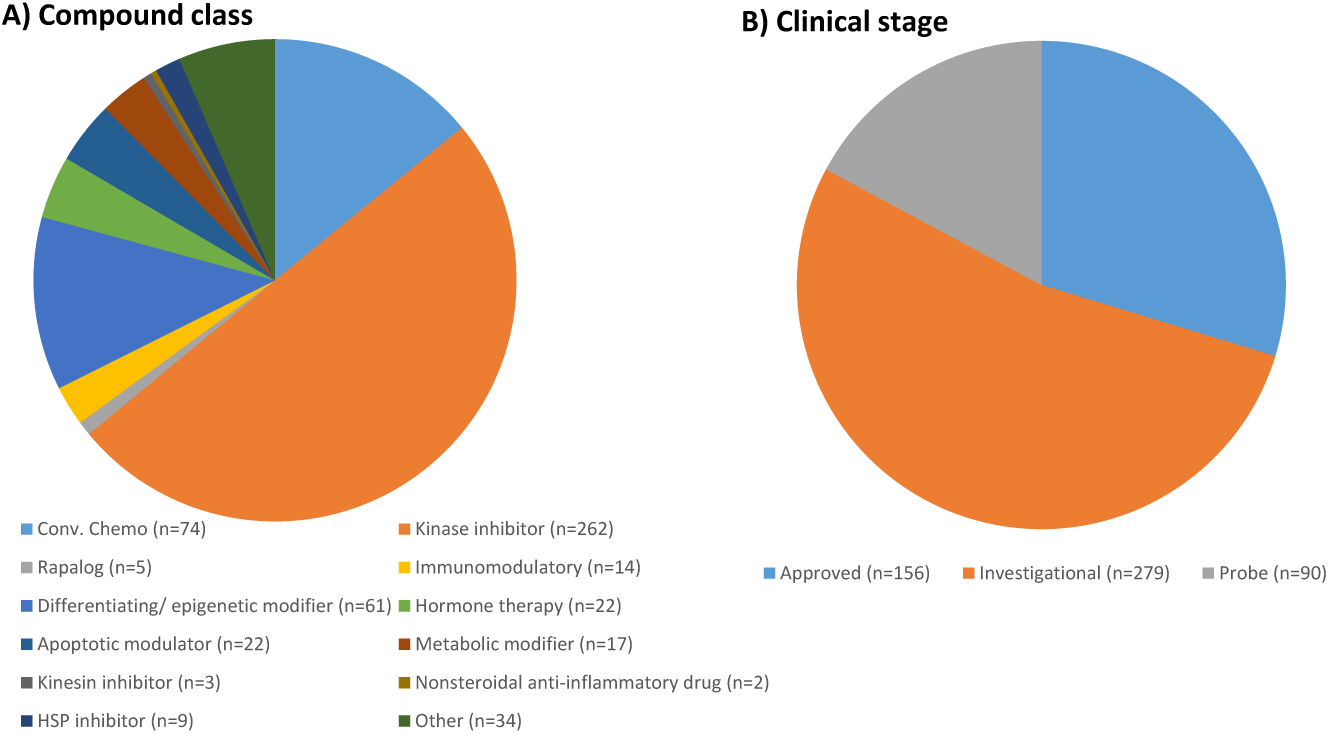
An overview of the FIMM oncology compound collection. The drug combination platform enables the testing of pairwise drug combinations from 525 small-molecular anticancer compounds that cover mainly kinase inhibitors and other signaling transduction modulators. A majority of the compounds are either FDA-approved or being evaluated in clinical trials at different stages.

1. The compounds are dissolved in DMSO except for 19 drugs (e.g. platinum drugs) with poor DMSO solubility or stability that instead are dissolved in water. All the 525 compounds are pre-drugged in five concentrations using eight 384-well plates ***(10)***.
2. The pre-drugged plates are stored in Storage Pods (Roylan Developments Ltd) under nitrogen gas at room temperature up to 1 month.
3. Quality control. As a regular quality check-up of our compound library, we are testing the full FO4B collection set with four assay-ready cell lines (DU4475, HDQ-P1, IGROV-1 and MOLM-13) every two months. Following the time-dependent reproducibility of the drug responses allows us to precisely detect any changes in the compound stability and activity.

### 2.3. Equipment

1. Labcyte Echo 550 acoustic dispenser for dispensing compounds in precise volume with high accuracy (2.5 nL).
2. MultiFlo FX Multi-Mode Dispenser with RAD module (BioTek) or Multidrop Combi Reagent Dispenser (Thermo Scientific) for dispensing growth media, CellTiter-Glo reagents and seeding cells.
3. Beckman Coulter Biomek FX^P^ for dispensing primary cells that tend to grow as aggregates.
4. PHERAstar FS (BMG Labtech) or Cytation 5 Cell Imaging (BioTek) multi-mode plate readers for CellTox Green (fluorescence) and CellTiter-Glo (luminescence) detection on 384-well plates.
5. 384-well tissue culture treated sterile assay plates (Corning).

### 2.4. Reagents

1. Cell media, serum and supplements recommended by cell line providers.
2. CellTox Green Cytotoxicity Assay for measuring dead cells (Promega).
3. CellTiter-Glo or CellTiter-Glo 2.0 Assay for detecting cell viability (Promega).

### 2.5. Software tools

Specific software tools are needed in the experimental design stage and in the data analysis stage. For the 384-well plate design, once the drugs and the concentration ranges are selected, we use the in-house cherry-picking program, FIMMcherry, to automatically generate the echo files needed for the Labcyte Access system. When the drug combination dose-response matrix data is ready, we then use the synergyfinder R package to score and visualize the drug interactions. The synergyfinder is also available as a web-application without the need to install the R environment.

## 3. Methods

The drug combination analysis pipeline starts from sample preparation and compound selection, based on which an automated plate design program called FIMMCherry is utilized. The phenotypic readout from the plate is then profiled including cell viability, cytotoxicity or other phenotypes. The resulting dose-response matrix data is analyzed with the synergyfinder R package for the detection of synergistic drug combinations (Fig. 2).

**Fig. 2.**
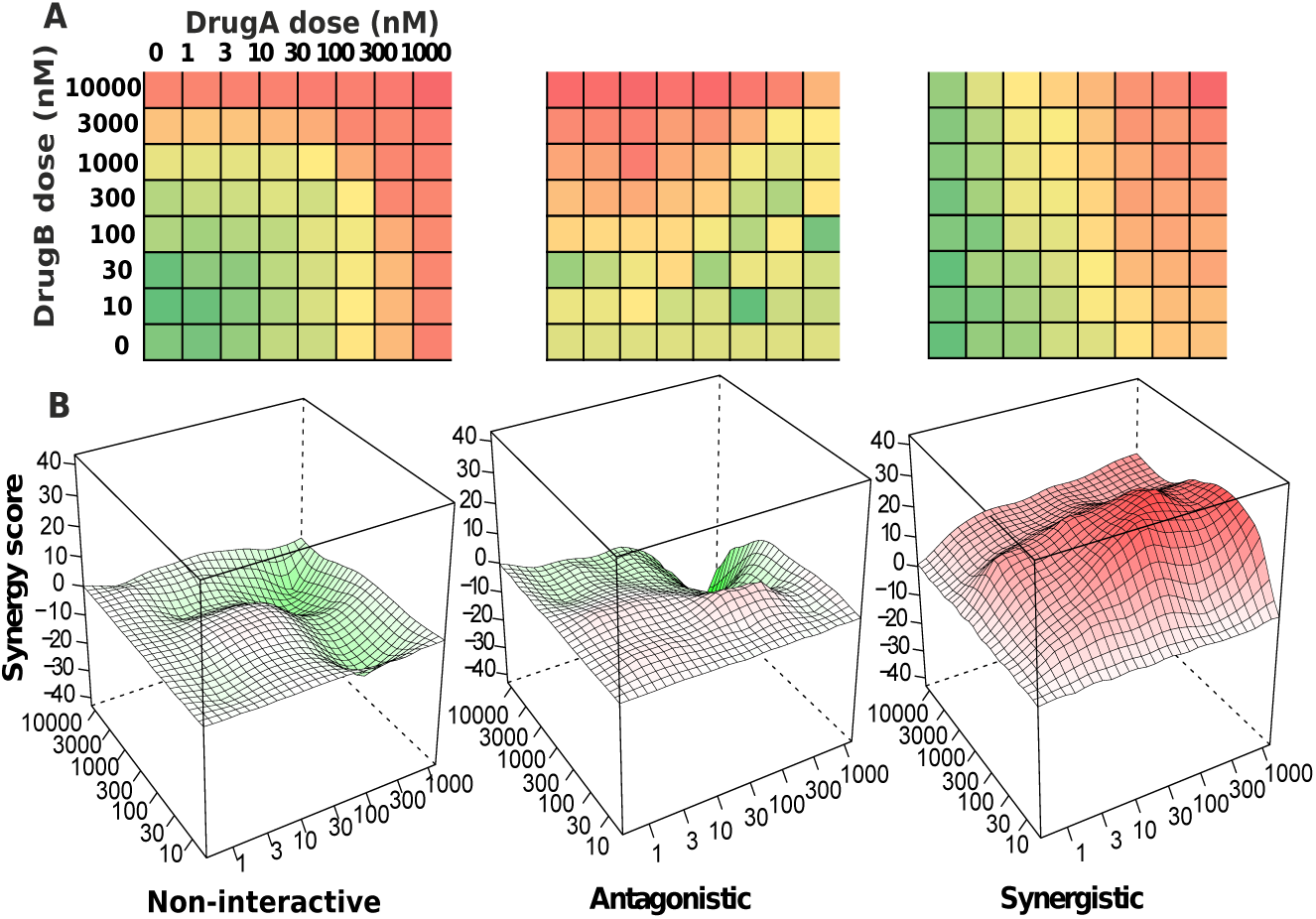
An overview of the drug combination data analysis. (A) A typical high-throughput drug combination screen utilizes a dose-response matrix design where all possible dose combinations for a drug pair can be tested. Colors in the dose-response matrices indicate different levels of phenotypic responses of the cancer cell. (B) Depending on the interaction pattern models derived from the dose-response matrices, a drug combination can be classified as non-interactive, antagonistic or synergistic.

### 3.1. Cell culture

1. Dissociation of cells by 0.05% trypsin-EDTA (Gibco) or HyQTase (HyClone) to single cell suspension.
2. Titrate cells to define optimal density within exponential growth (log phase). Cell seeding in 2-fold serial dilution starting from 16,000 cells/well on 384-well plates. For most cell lines, the optimal cell number is in the range of 500-2,000 cells/well.
3. Cell toxicity and viability detection after 72 h of incubation using CellTox Green and CellTiter-Glo reagents (Promega).
4. For microenvironment control to minimize edge effect and keep concentrations of solutions constant we are using MicroClime Environmental Lids (Labcyte).

### 3.2. Drug combination plate design

We utilize a combination plate layout where six compound pairs can be accommodated on one 384-well plate. A given pair of drugs is combined in a series of one blank and seven half-log dilution concentrations, resulting in an 8 × 8 dose matrix. To be able to transfer the compounds according to this matrix format, a pick list defining the source and destination plate locations and transfer volumes for the compounds is needed. An in-house program, called FIMMCherry, has been developed to automatically generate these rather complex pick lists effortlessly.

FIMMCherry is a desktop GUI application, which is developed using Python (https://www.python.org/) and Qt application development framework (https://www.qt.io/). The integration of Python and Qt allows FIMMCherry to run on all the major computer platforms including Windows, Linux and Mac OS X. Two tab-delimited text files are needed as input:

1. A source plate file provides information of the compound stocks (compound identification, available concentration ranges, source plate identification and well identification);
2. A drug combination file containing the selected compounds.

After loading the input files, FIMMCherry will show the layout of the plates accordingly (Fig. 3). A pick list that is compatible with the Labcyte Echo dispenser is then provided for compound dispensing.

**Fig. 3.**
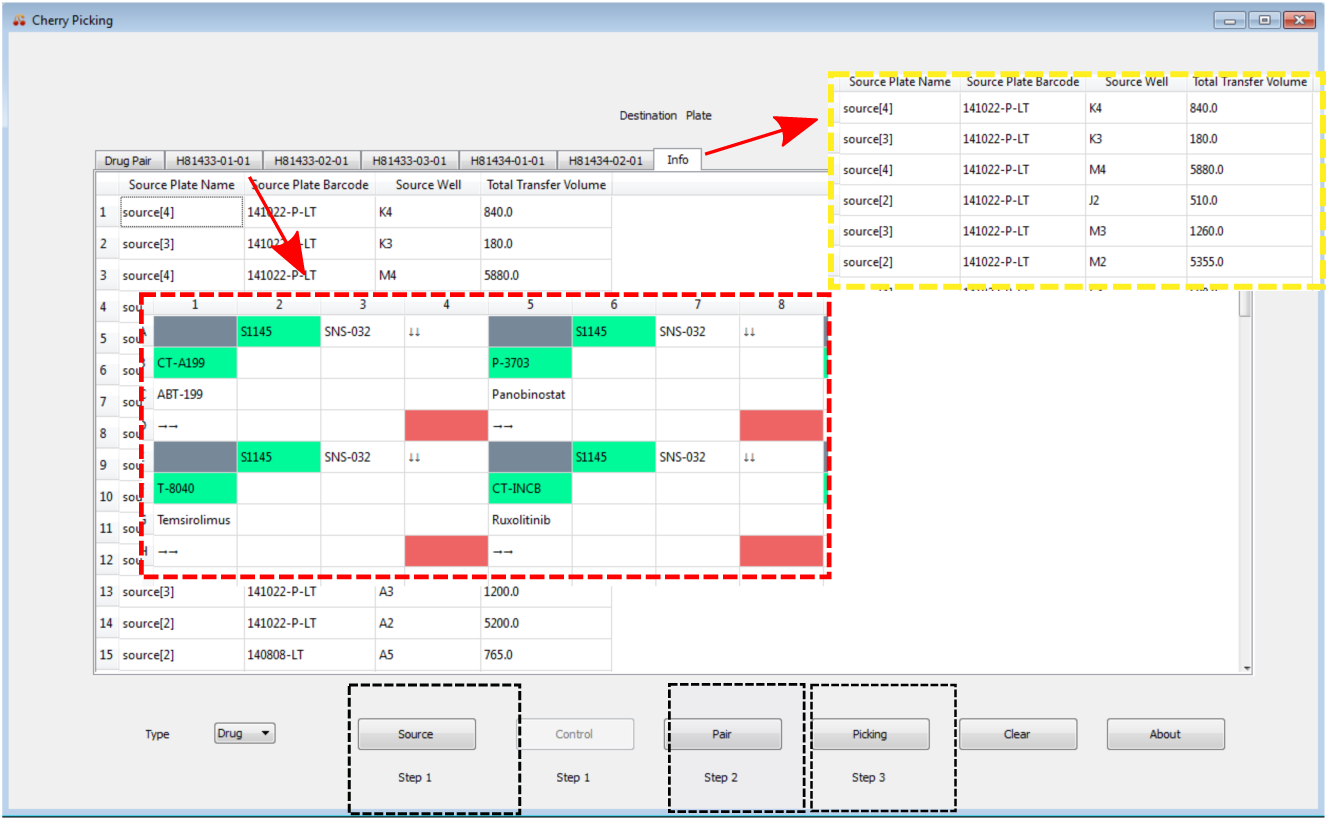
Drug combination plate design using FIMMCherry. The graphical user interface contains a virtual plate enabling an interactive way of design. After loading the input files including the source, the control and drug pair information (the black inset boxes), the selected drug combinations and their dose ranges will be listed in the ‘Drug Pair’ tab, for which an echo file will be generated for acoustic dispensing. Each plate can be visualized in a separate tab, named by their plate identifiers (the red inset box). The ‘Info’ tab shows the liquids consumption in the source plates (the yellow inset box).

### 3.3. Phenotypic readouts

1. Transfer 5 uL of media with CellTox Green Cytotoxicity reagent in 384-well pre-drugged plate.
2. Shake the plates on the plate shaker at 450 rpm for 5 min for proper drugs dissolving.
3. Transfer a single cell suspension in 20 uL of media to 384-well plate. Final dilution of CellTox Green reagent should be 1:2,000 in 25 uL.
4. Incubate cells on the plates during 72 h.
5. Shake the plates on the plate shaker at 500 rpm for 30 s. Read fluorescence on the plates by plate reader for CellTox Green Cytotoxicity detection.
6. Transfer 25 uL of CellTiter-Glo reagent.
7. Shake the plates on the plate shaker at 450 rpm for 5 min.
8. Read luminescence on the plates for detecting cells viability using CellTiter-Glo assay.

### 3.4. Synergy scoring

#### 1. Installation of the synergyfinder R package

After downloading and installing R (https://www.R-project.org) and Bioconductor (https://www.bioconductor.org/), the synergyfinder package can be installed automatically by typing in the R console as below:

> source(“https://www.bioconductor.org/biocLite.R”)
> biocLite(“synergyfinder”)

#### 2. Input data

A single csv file that describes a drug combination dataset is provided as the input. The csv file is in a list format and must contain the following columns:

• BlockID: the identifier for a drug combination. If multiple drug combinations are present, e.g. in the standard 384-well plate where 6 drug combinations are fitted, then the identifiers for each of them must be unique.
• Row and Col: the row and column indexes for each well in the plate.
• DrugCol: the name of the drug on the columns in a dose-response matrix
• DrugRow: the name of the drug on the rows in a dose-response matrix
• ConcCol and ConcRow: the concentrations of the column drugs and row drugs in a combination
• ConcUnit: the unit of concentrations. It is typically nM or μM.
• Response: the effect of drug combinations at the concentrations specified by ConcCol and ConcRow. The effect must be normalized to %inhibition based on the positive and negative controls. For a well-controlled experiment, the range of the response values is expected from 0 to 100. However, missing values or extreme values are allowed. For input data where the drug effect is represented as %viability, the program will internally convert it to %inhibition value by 100-%viability.

We provide an example input data in the R package, which is extracted from a recent drug combination screening for the treatment of diffuse large B-cell lymphoma (DLBCL) ***(7)***. The example input data contains two representative drug combinations (ibrutinib & ispinesib and ibrutinib & canertinib) for which the %viability of a cell line TMD8 was assayed using a 6 by 6 dose matrix design. The example data in the required list format can be loaded and reshaped to a dose-response matrix format for further analysis by typing:

> data(“mathews_screening_data”)
> dose.response.mat <- ReshapeData(mathews_screening_data, data.type = “viability”)

The ‘data.type’ parameter specifies the type of drug response, which can be either ‘viability’ or ‘inhibition’. We will use these example data to illustrate the main functions of synergyfinder below.

More documentation of the input and output parameters for each function can be accessed by typing:

> help(‘ReshapeData’)

#### 3. Input data visualization

The input data can be visualized using the function PlotDoseResponse by typing:

> PlotDoseResponse(dose.response.mat)

The function fits a four-parameter log-logistic model to generate the dose-response curves for the single drugs based on the first row and first column of the dose-response matrix. The drug combination responses are also plotted as heatmaps, from which one can assess the therapeutic significance of the combination, e.g. by identifying the concentrations at which the drug combination can lead to a maximal effect on cancer inhibition (Fig. 4). The PlotDoseResponse function also provides a high-resolution pdf file by adding the ‘save.file’ parameter:

> PlotDoseResponse(dose.response.mat, save.file = TRUE)

**Fig.4.**
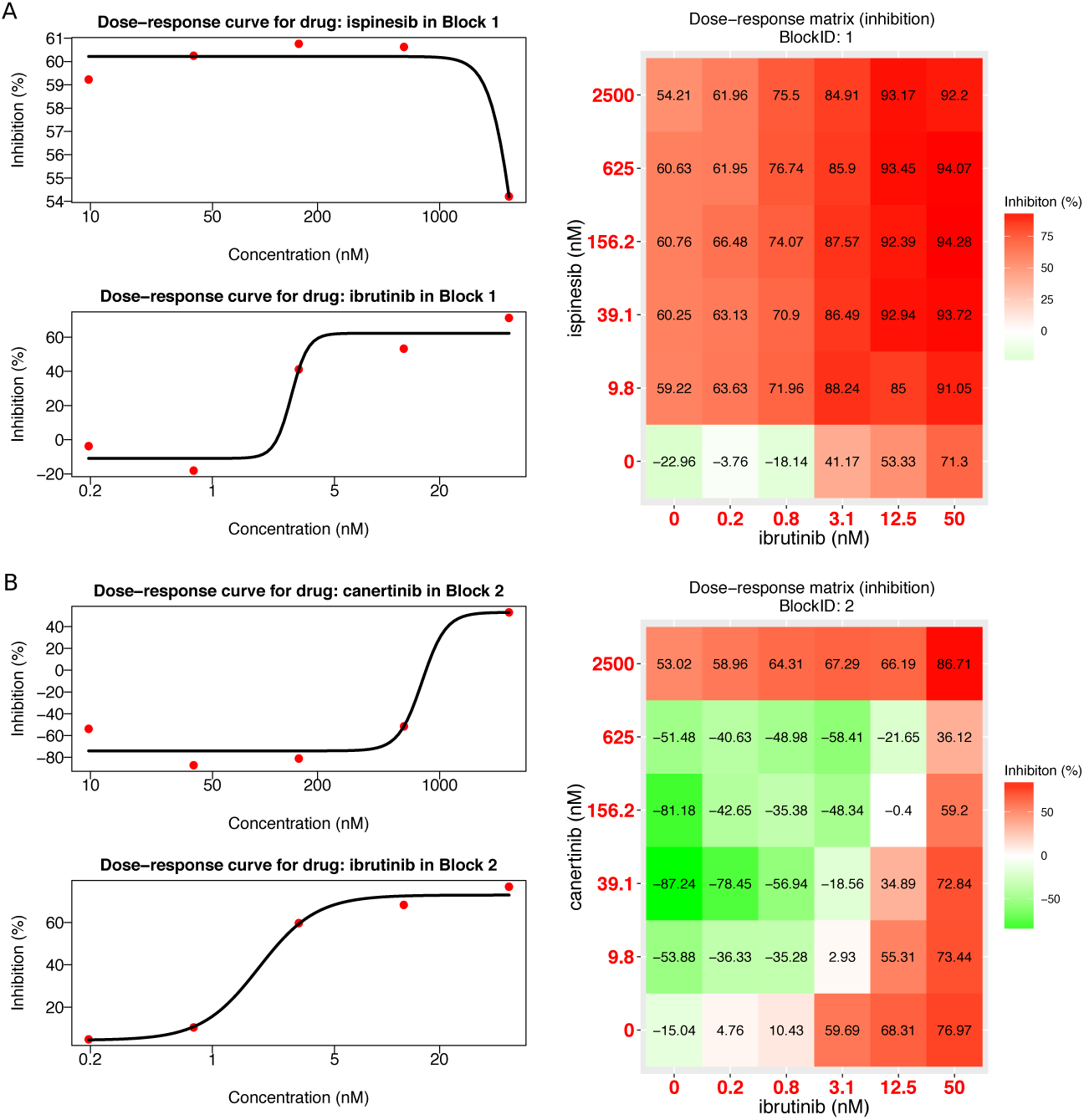
Plots for single drug dose-response curves and drug combination dose-response matrices. (A) The ibrutinib and ispinesib combination. (B) The ibrutinib and canertinib combination. Left panel: single drug dose-response curves fitted with the commonly-used 4-parameter log-logistic (4PL) function. Right panel: the raw dose-response matrix data is visualized as a heatmap.

The pdf file will be saved under the current work directory with the syntax: “drug1.drug2.dose.response.blockID.pdf.

#### 4. Drug synergy scoring

The current synergyfinder package provides the synergy scores of four major reference models, including HSA, Loewe, Bliss and ZIP. Let’s consider a drug combination experiment where drug 1 at dose *x*_1_ is combined with drug 2 at dose *x*_2_. The effect of such a combination is *y*_*c*_ as compared to the monotherapy effect *y*_1_(*x*_1_) and *y*_2_(*x*_2_). To be able to quantify the degree of drug interactions, one needs to determine the deviation of *y*_c_ from the expected effect *y*_e_ of non-interaction, which is calculated in different ways with the reference models.

• HSA: *y*_e_ is the effect of the highest monotherapy effect, i.e. *y*_e_ = *max* (*y*_1_, *y*_2_).
• Loewe: *y*_e_ is the effect as if a drug is combined with itself, i.e. *y*_e_ = *y*_1_(*x*_1_+*x*_2_) = *y*_2_(*x*_1_+*x*_2_)
• Bliss: *y*_e_ is the effect as if the two drugs are acting independently on the phenotype, i.e. *y*_e_=*y*_1_+*y*_2_-*y*_1_*y*_2_
• ZIP: *y*_e_ is the effect as if the two drugs do not potentiate each other, i.e. both the assumptions of the Loewe model and the Bliss model are met.

Once *y*_e_ can be determined, the synergy score can be calculated as the difference between the observed effect *y*_c_ and the expected effect *y*_e_. Depending on whether *y*_c_ > *y*_e_ or *y*_c_ < *y*_e_ the drug combination can be classified as synergistic or antagonist, respectively. Furthermore, as the input data has been normalized as %inhibition values then the synergy score can be directly interpreted as the proportion of cellular responses that can be attributed to the drug interactions.

For a given dose-response matrix, one need to first choose what reference model to use and then apply the CalculateSynergy function to calculate the corresponding synergy score at each dose combination. For example, the ZIP-based synergy score for the example data can be obtained by typing:

> synergy.score <- CalculateSynergy(data = dose.response.mat, method = “ZIP”, correction = TRUE)

For assessing the synergy scores with the other reference models, one needs to change the ‘method’ parameter to ‘HSA’, ‘Loewe’ or ‘Bliss’. The ‘correction’ parameter specifies if a baseline correction is applied on the raw dose-response data or not. The baseline correction utilizes the average of the minimum responses of the two single drugs as a baseline response to correct the negative response values. The output ‘synergy.score’ contains a score matrix of the same size to facilitate a dose-level evaluation of drug synergy as well as a direct comparison of the synergy scores between two reference models.

#### 5. The drug interaction landscape

The synergy scores are calculated across all the tested concentration combinations, which can be straightforwardly visualized as either a two-dimensional or a three-dimensional interaction surface over the dose matrix. The landscape of such a drug interaction scoring is very informative when identifying the specific dose regions where a synergistic or antagonistic drug interaction occurs. The height of the 3D drug interaction landscape is normalized as the % inhibition effect to facilitate a direct comparison of the degrees of interaction among multiple drug combinations. In addition, a summarized synergy score is provided by averaging over the whole dose-response matrix. To visualize the drug interaction landscape, one can utilize the PlotSynergy function as below (Fig. 5):

> PlotSynergy(synergy.score, type = “all”, save.file = TRUE)

**Fig.5.**
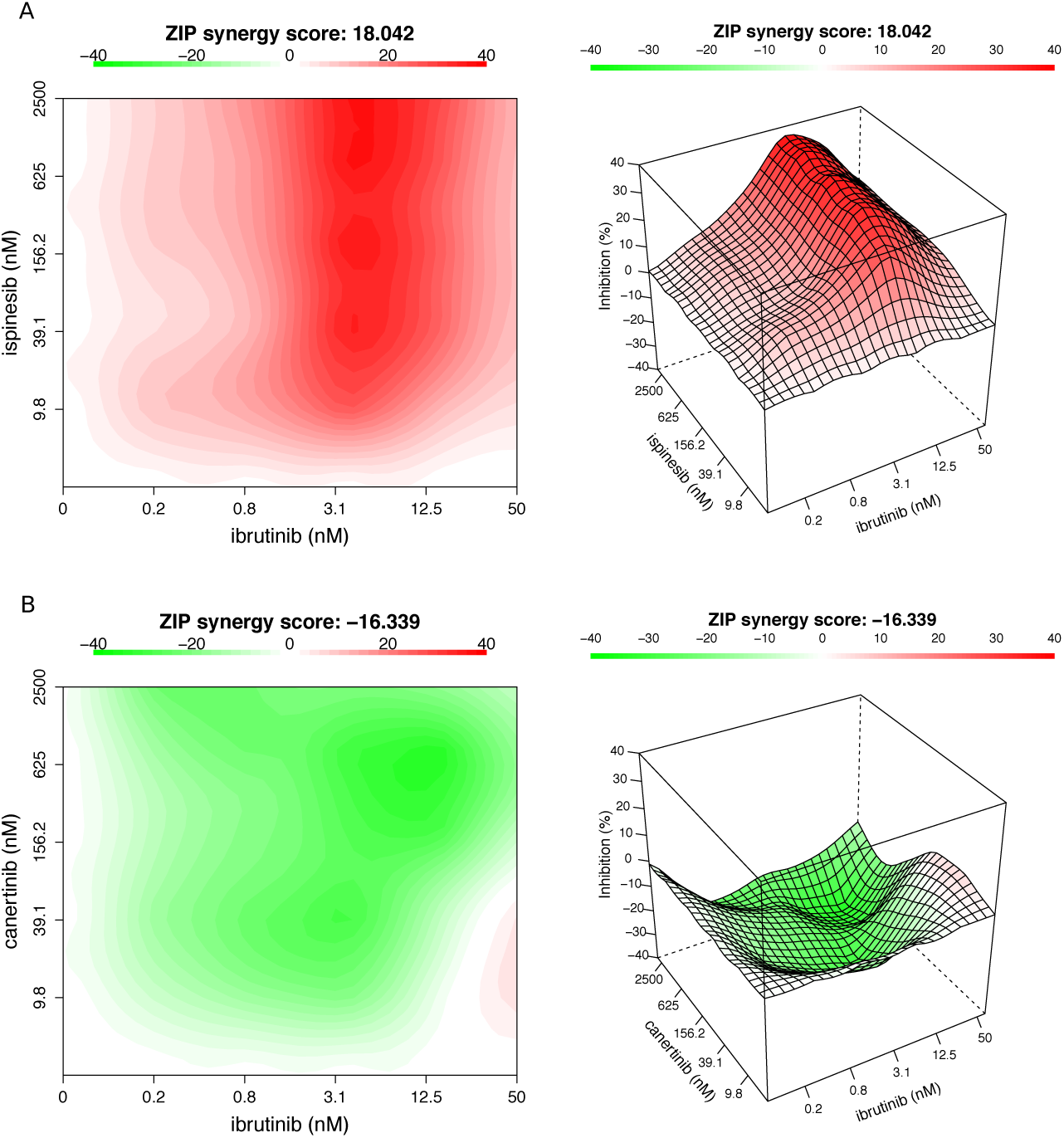
The drug interaction landscapes based on the ZIP model. (A) The ibrutinib and ispinesib combination. (B) The ibrutinib and canertinib combination.

The ‘type’ parameter specifies the visualization type of the interaction surface as 2D, 3D or both.

## 4. Notes

To identify potential drug combinations in preclinical settings both appropriate experimental techniques and computation methods are required. However, many of the drug interaction scoring methods are focused on a theoretical advance in mathematical modeling, while their corresponding implementation tools or source codes are seldom made easily accessible, which hinders their application in the analysis of concurrent drug combination data ***(26–27)***. We provide the R package synergyfinder to calculate the drug synergy scores using four different reference models, acknowledging the fact that the standardization of drug combination data analysis remains an open question ***(28)***. The users are therefore advised to apply all the models for their data and report a drug combination that can show a detectable level of synergy scores irrespective of the model in selection. Further, a strong synergy in a drug combination, as revealed using the synergy landscape analysis might not be sufficient to warrant the next level confirmatory analysis if the synergy does not lead to sufficient overall responses. Therefore, the synergy scoring is always advised to be combined with the raw dose-response matrix data visualized in Fig. 4 to provide an overview of the extra benefits of drug combinations compared to single drugs. The synergyfinder package will be continuously updated for including more rigorous analyses such as statistical significance, effect size and noise detection.

Availability: The source code for the FIMMCherry program is available at github (https://github.com/hly89/FIMMCherry). The synergyfinder R package for drug combination data analysis is available at CRAN and Bioconductor.

## Acknowledgements

This work was supported by the Academy of Finland (grants 272437, 269862, 279163, 295504 and 292611 for TA, 272577 and 277293 for KW); the Integrative Life Science Doctoral Program at the University of Helsinki (LH), the Sigrid Jusélius Foundation (KW) and the Cancer Society of Finland (JT, TA and KW). This project has received funding from the European Union’s Horizon 2020 research and innovation program 2014–2020 under Grant Agreement No 634143 (MedBioinformatics).

